# Mad Hatter correctly annotates 98% of small molecule tandem mass spectra searching in PubChem

**DOI:** 10.1101/2022.12.07.519436

**Authors:** Martin A. Hoffmann, Fleming Kretschmer, Marcus Ludwig, Sebastian Böcker

## Abstract

Metabolites provide a direct functional signature of cellular state. Untargeted metabolomics usually relies on mass spectrometry, a technology capable of detecting thousands of compounds in a biological sample. Metabolite annotation is executed using tandem mass spectrometry. Spectral library search is far from comprehensive, and numerous compounds remain unannotated. So-called *in silico* methods allow us to overcome the restrictions of spectral libraries, by searching in much larger molecular structure databases. Yet, after more than a decade of method development, *in silico* methods still do not reach correct annotation rates that users would wish for. Here, we present a novel computational method called Mad Hatter for this task. Mad Hatter combines CSI:FingerID results with information from the searched structure database via a metascore. Compound information includes the melting point, and the number words in the compound description starting with the letter ‘u’. We then show that Mad Hatter reaches a stunning 97.6% correct annotations when searching PubChem, one of the largest and most comprehensive molecular structure databases. Finally, we explain what evaluation glitches were necessary for Mad Hatter to reach this annotation level, what is wrong with similar metascores in general, and why metascores may screw up not only method evaluations but also the analysis of biological experiments.

## Introduction

Metabolomics, the comprehensive analysis of metabolites in a biological organism or system, provides a functional readout of its physiological state. Metabolites cover literally thousands of compound classes; their structural diversity is much larger than for biopolymers such as proteins. Many metabolite structures remain to be discovered, in particular for non-model organisms and microbial communities; but even the human metabolome still holds surprises, such as the 2018 discovery of an antiviral metabolite^1^. Non-targeted metabolomics experiments usually rely on mass spectrometry (MS), where tandem MS (MS/MS) is used for structural annotations. This experimental setup can detect thousands of molecules in a biological sample.

Spectral libraries store MS/MS spectra of “known compounds”, meaning that the identity of the compound behind each spectrum is known. Large and comprehensive libraries are needed for annotating MS/MS spectra via library search. Importantly, not the number of spectra but the number of compounds is driving the amount of information present in a spectral library. Unfortunately, spectral libraries are not only substantially smaller than molecular structure databases, they also grow at a much slower pace: The vast majority of MS/MS data in a spectral library are *reference measurements* from synthetic standards. Again in the vast majority of cases, those standards are commercial compounds ordered from a vendor. When a “new spectral library” is measured in a lab, it usually contains almost exclusively compounds that are already available in other libraries. Hence, the number of MS/MS spectra is growing, but not the number of compounds; and as noted above, more spectra add little information. This is because standards become more and more expensive, the further we “leave the beaten path” of synthetic chemistry.

In the last decade, development of computational methods for analyzing metabolite MS and MS/MS data has gained momentum^2–8^. Six CASMI contests (Critical Assessment of Small Molecule Identification) were conducted from 2012 to 2022; these are blind competitions evaluating computational methods, in particular for searching in a molecular structure database using a query MS/MS spectrum^9–11^. This particular task is a striking example of the differences between metabolomics and shotgun proteomics: Even the simplest approaches for searching a peptide MS/MS spectrum in a structure (that is, peptide sequence) database result in good annotation rates; in contrast, despite of more than a decade of research, development of so-called *“in silico* methods” for searching in molecular structure databases remains a challenging problem in metabolomics.

In the following, we concentrate on metabolomics and the task of metabolite annotation. Clearly, our method can also be used in other areas such as environmental research, where compound annotation from MS data is sought.

## Results

Here, we introduce Mad Hatter as a novel method for small molecule annotation. MS/MS data of the query compound is matched against a structure database using a metascore: Mad Hatter combines CSI:FingerID search results with information available from the searched molecular structure database. Used features include the melting point of the compound, its PubChem^12^ CID number modulo 42, and the number of occurrences of the sequence of letters “Was it a cat I saw” in a compound’s PubChem record. See Table 1 and Methods section for details. To combine features and the CSI:FingerID score, we use a state-of-the-art machine learning model, namely, a deep neural network with two dense hidden layers of 256 neurons each. Notably, Mad Hatter does not use features previously suggested for metascores, such as citation counts, production volumes, or the number of structure databases a compound is contained in.

**Table 1.** Features used by Mad Hatter. Features are read from the JSON file of a PubChem entry using field “name”. In addition to the information from PubChem, Mad Hatter uses the CSI:FingerID score as a feature.

| Used field | Feature description |
| --- | --- |
| RecordNumber | PubChem CID modulo 42 |
| RecordTitle | Record name length |
| References, XLogP | Number of information sources × XLogP value, absolute value |
| Complexity | Structure complexity (numerical value) |
| Melting Point | Melting point (numerical value) |
| Stability/Shelf Life | Number of vowels in the shelf life description |
| Synonyms | Number of consonants in a compound’s synonym list |
| Record Description | Number of occurrences of word “DNA” in compound description |
| full record | Number of words starting with letter ‘U’ (case insensitive) in a compound’s record |
| full record | Number of occurrences of “WAS IT A CAT I SAW” in a compound’s record (case insensitive, blanks ignored) |

We evaluate Mad Hatter using the well-known test dataset from CASMI 2016 (positive ion mode data). Mad Hatter correctly annotated 97.6% of query mass spectra when searching in PubChem, a structure database containing about 100 million compounds. To the best of our knowledge, this is the highest annotation rate ever reported for any in *silico* tool. We infer that Mad Hatter is currently the best tool for compound annotation. We also trained an alternative, less powerful version of Mad Hatter using a smaller, less powerful neural network with only one hidden layer with 12 neurons. Using this smaller, less powerful neural network we still reached 79.5% correct annotations.

Measuring MS/MS spectra needs a lot of time and effort. We found that we can reach 72.4% correct annotations *without* MS/MS data, just using the precursor *m/z* and the metascore (small neural network). This is also possible if we replace our MS/MS spectra by random spectra (71.7% correct annotations) or by empty spectra (71.0% correct annotations). These numbers are on par with those of any *in silico* method that participated in the original CASMI 2016 contest^9^. Hence, we can basically replace all parts of the mass spectrometry instruments responsible for measuring MS/MS data by a simple random number generator, or by a device that generates zeroes. We conjecture that this simple trick may save us considerable amount of money and time in the future.

## Methods

### Datasets and molecular structure database

Training and test dataset were downloaded from the CASMI 2016 web page at http://www.casmi-contest.org/2016/challenges-cat2+3.shtml. We restricted ourselves to the data measured in positive ion mode. All evaluations could easily be replicated for negative ion mode data. The *training dataset* consists of 254 instances with 242 unique molecular formulas. Each instance is a merged experimental MS/MS spectrum from one query compound, and is called a “challenge” in CASMI nomenclature. The *test dataset* consists of 127 instances with 125 unique molecular formulas. In four instances of the training dataset, respective molecular formulas were also present in the test dataset. These instances were removed from the training dataset, to guarantee structure disjointness between training and test dataset. This leaves 250 instances in the training dataset. Molecular structures were standardized using the PubChem standardization procedure^12^, using the web service at https://pubchem.ncbi.nlm.nih.gov/rest/pug/.

Molecular structures from the *PubChem structure database* were downloaded on 16 January 2019 and contain 97,168,905 compounds and 77,153,182 unique covalently bonded structures with mass up to 2,000 Da. JSON files containing information on those PubChem records were downloaded on 04 July 2022.

### Features and samples

Features were read from the downloaded JSON files. In general, features are extracted using the field names given in Table 1. The number of information sources is the amount of key-value pairs in the field `Reference`. For melting point, the first entry in the field `Melting Point` is taken. If no melting point is provided, we enter 0 K. The number of consonants in a compound’s synonyms is taken by iterating over all key-value pairs of the Synonyms field. To count the number of times the record `Descript` ion contains the string “DNA”, we actually count how often the string DNA can be constructed if we read the field from left to right. This is done case-insensitive. For example, “dragon” contains the string “DNA” zero times, “Modena” contains it once, and “Diana’s donat” contains it twice. The same is done searching the full record for “was it a cat I saw”, ignoring blanks. The `CSI score` was computed using SIRIUS 5.6.0 and CSI:FingerID 2.7.0, using structure disjoint cross-validation.

Each sample consists of a CASMI instance (query MS/MS) and a molecular structure candidate. As detailed below, we retrieve candidates from PubChem using the molecular formula of the query compound. Features were determined using the candidate structure; only the CSI:FingerID score feature considers both the candidate structure and the query MS/MS. Samples were labeled “correct” if a candidate’s structure and the correct query structure are identical, and “incorrect” otherwise. The training dataset consists of 250 samples labeled “correct” and 645,880 samples labeled “incorrect”. The test dataset consists of 127 samples labeled “correct” and 244,356 samples labeled “incorrect”.

### Neural network architecture and training

For the *large model,* we used a network architecture consisting of two fully connected hidden layers with 256 neurons each. Both hidden layers use the rectified linear activation function (ReLU). The output layer uses a sigmoid activation function and as such, returns the estimated probability of the input belonging to the positive class. For the *small model,* we used a network architecture consisting of one fully connected hidden layer, with 12 neurons. The hidden layer again uses the ReLU activation function. We used the same output layer as in the large architecture, but add a 30% dropout^13^ between the hidden layer and the output layer. Both architectures were trained with a batch size of 32,000 and for 5,000 epochs. We used the binary cross-entropy as loss function and Adam as the optimizer^14^. Features were standardized by setting their mean to zero, and standard deviation to one. Training and standardization was carried out using TensorFlow^15^.

### Evaluation

We trained and evaluated both models on the test dataset. For each of the 127 instances (queries), we assume that the correct molecular formula of the compound is known, as explained in [16, 17]. Therefore, for each instance the pool of candidates consists of those PubChem structures with the correct molecular formula. For every candidate compound, we calculated features as described earlier and predicted the probability of that compound being labeled “correct” using our trained model. The candidate compound with the highest probability is then returned as the annotation for that instance.

As noted above, we employ CSI:FingerID ensuring structure disjoint cross-validation: For each query compound, we choose the cross-validation CSI:FingerID model so that the compound structure is not in the training data. See [16, 17] for details.

### Random mass spectra and empty mass spectra

To generate an *empty MS/MS spectrum,* we inserted a precursor peak with intensity 100% into the otherwise empty MS/MS spectrum. For that, we used the *m/z* of the MS1 peak with the highest intensity in a 10 ppm window around the compound *m/z.* To generate a *random MS/MS spectrum*, we first inserted the precursor peak with *m/z* of *M* as described above. Then, we added 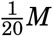 random peak *m/z* uniformly distributed in [0,M]. Intensities were drawn uniformly from [0,1]. Evaluations were performed using the Mad Hatter small model trained on the training dataset. Again, we employ the CSI:FingerID model from cross-validation that has not seen the compound structure.

## Remorse

It might not have escaped the reader’s attention that we do not seriously propose to use Mad Hatter for any practical compound annotation. In fact, besides choosing outrageous features for Mad Hatter, we deliberately introduced a series of blunders in our evaluations that allowed us to reach the reported annotation rates. We start off with describing the blunders, and discuss the outrageous features in the next section. Unfortunately, the mentioned blunders can regularly be found in other scientific publications that present novel *in silico* methods; since this is not blame- and-shame, we will refrain from explicitly pointing out those papers. We argue that it is rather editors and reviewers than authors who are responsible for the current unpleasant situation. The blunders we deliberately made in the development and evaluation of Mad Hatter may serve as a guide to editors and reviewers for spotting similar issues “in the wild”.

Early publications^16,18,19^ went through a great deal of effort to demonstrate and evaluate not only the power but also the *limitations* of the presented methods. In contrast, it lately became a habit to demonstrate that a new method strongly outcompetes all others in some evaluation. It is sad to say that those claims are often pulverized later in blind CASMI challenges. One way to achieve the desired outcome is to carefully select our competitors; leave out those that are hard to beat. Another possibility is to evaluate other methods ourselves but not invest too much time into this part of the evaluation; nobody will mind the inexplicable performance drop of the other method unless we explicitly mention it. A third possibility is to restrict reported results to one evaluation dataset where the desired outcome is reached. Finally, we may simply present results as if our method was the first to ever deal with the presented problem, and simply ignore the presence of other methods; this is the approach we chose for Mad Hatter. Yet, the most common trick to make our method shine, is: Evaluate on data our method already knows from training. This is commonly referred to as “data leakage”.

### Do not evaluate on your training data

For the Mad Hatter evaluation, the first blunder to notice is that we do not differentiate between training set and the dataset used for evaluation. Consequently, the machine learning model is rewarded for memorizing instead of learning. This must be regarded a show stopper for a method publication, as all numbers reported are basically useless. In fact, CASMI 2016 offers two independent datasets (train and test) that can be used for exactly this purpose. If we repeat the above evaluation but this time, make a clear distinction between training and test set, correct annotations drop to 61.4% using the large neural network. This might still be regarded as a good annotation rate and might even get published. Yet, the fact that we observe such a *massive drop in annotation rates* must be regarded as a red flag that our machine learning model is overfitting. This red flag shines even brighter if we compare to the annotation rate of CSI:FingerID *without* any metascore addition: It turns out that CSI:FingerID reaches 49.6% correct annotation in a structure-disjoint evaluation. Somehow, we forgot to report this important baseline number in the Results section.

In fact, we do not even have to know those numbers from evaluation, to suspect that something fishy is going on: We have trained a neural network with 68.3k parameters on a training dataset containing 250 queries, corresponding to 250 positive samples (see Methods). Even tricks such as regularization, dropout^13^, or batch normalization^20^ — which we deliberately did not use — could not prevent that the resulting model is overfitting to the training data. If the number of training samples is small, then the use of a large “vanilla” deep neural network (also known as multilayer perceptron) should raise some doubt. During the last years, numerous network architectures were developed that substantially reduced the number of parameters in the model, in order to improve its generalization capabilities: Convolutional neural networks for image recognition are a prominent example. Yet, unless there is a convincing reason why such a network architecture should be applied for the problem at hand, we cannot simply grab a random architecture and hope that it will improve our results! Stating that a particular model architecture is “state-of-the-art” in machine learning, does not mean it is reasonable to be applied. Machine learning cannot do magic.

For generalizability, less is often more: As reported above, we also trained a sparser geometry of the neural network with only one hidden layer of 12 neurons. This implies that we have substantially fewer parameters in our model. In contrast to our claims in the results section, this does not mean that the model is less powerful. Rather, there is a good chance that the smaller model will perform better on any independent data. We deliberately made it even harder for the small model to “learn”, using 30% dropout when training the network^13^. This means that 30% of the neurons in the hidden layer are randomly removed before forward and backpropagation is used to update edge weights. Doing so considerably worsened the performance of the model when trained on the evaluation data (memorizing) but simultaneously, improved the performance when trained on the training data and evaluated on the evaluation data (learning). For this more sensible machine learning model, we reach 79.5% correct annotations when training and evaluating on the test dataset, and 76.4% when training on the training data and evaluating on the evaluation data.

### Do not evaluate on your training data, reloaded

Unfortunately, separation of training and test set is not as easy as one may think. For an *in silico* method, saying “we used dataset A for training and dataset B for evaluation” is insufficient. The first thing to notice is that the same spectra are often found in more than one spectral library, and we have to ensure that we do not train on the same data we evaluate on. A prominent example are the data from the CASMI 2016 contest, that have been repeatedly used for method evaluations. These data were later included in spectral libraries such as GNPS (Global Natural Products Social) and MoNA (MassBank of North America). Hence, a method trained on GNPS and evaluated on CASMI 2016 has seen all the data, and the reported numbers are meaningless for an *in silico* method.

Yet, even a *spectrum-disjoint* evaluation is insufficient: If the same compound is measured twice for two libraries, then the resulting spectra will be extremely similar. In fact, library search in public and commercial spectral libraries is based on this observation. Hence, we must ensure that the same compound is never found in training and test data, to avoid that we are basically evaluating spectral library search — which we already know to work rather well. Unfortunately, the rabbit hole goes deeper: This *compound-disjoint evaluation* is still insufficient. Some compounds (say, L- and D-threose, threose and erythrose) may just differ in their stereochemistry. Yet, stereochemistry has only a minor impact on fragmentation of a compound. Hence, evaluations of *in silico* methods ignore stereochemistry, and evaluate search results at the level of molecular structures. (At present, an *in silico* method evaluation claiming to differentiate L/D-threose and L/D-erythrose based on MS/MS data, must not be taken seriously.) To this end, we have to treat all those compounds as the same *structure*, and ensure that our evaluation is *structure-disjoint*. This is the minimum requirement for an *in silico* evaluation to be reasonable.

There are situations that require an even stricter separation of training and test data: For example, the second evaluation of Mad Hatter was executed *molecular formula-disjoint,* meaning that compounds with the same molecular formula must not appear in training and test data. This implies that the evaluation is also structure-disjoint. The stricter molecular formula-disjoint evaluation is advisable in cases where one suspects that the molecular formula is already giving away too much information to a machine learning model. Finally, *scaffold-disjoint* evaluations ensure that compounds with the same scaffold must not appear in training and test data; such evaluations are common in virtual screening. From our experience, we can say that a scaffold-disjoint evaluation is not mandatory for *in silico* methods that analyze MS/MS data. Minor side group changes to a scaffold sometimes strongly affect fragmentation and sometimes not, making it impossible for machine learning to simply memorize spectra. Yet, for methods that consider fragmentation through Electron Ionization (EI), a scaffold-disjoint evaluation is mandatory! Here, each compound is found many times in the library with different derivatizations, as these derivatizations are part of the experimental protocol. If we evaluate the machine learning model on a compound for which ten derivatives are present in the training data, chances are high that the model can spot some similarity to one of the derivatives and basically falls back to memorizing.

Some methods employ machine learning without clearly stating so: Assume that a method adds the scores of two *in silico* tools. Clearly, we require a multiplier for one of the scores, to make them comparable. Yet, how do we find this multiplier? In fact, it is not too complicated to come up with a set of feature multipliers for Mad Hatter, avoiding the machine learning part of the manuscript. Yet, what we are building — manually or automated — is (highly similar to) a linear classifier from machine learning. Hence, our evaluations must be carried out structure-disjoint (for MS/MS data) or scaffold-disjoint (for EI data). Results that fail to do so are basically meaningless.

## Metascores

Unlike many publications from the last years, the resulting procedures for training and evaluation of Mad Hatter now follow established standards in machine learning. In fact, we went slightly beyond those standards by using a molecular formula-disjoint evaluation setup. Yet, we have used features that are quite obviously nonsense and should not be used for a metascore. Even less so, they should allow us to annotate compounds at the rates reported above! Recall that in a molecular formula-disjoint evaluation, Mad Hatter reached 76.4% correct annotations when searching PubChem. How is this possible? Thing is, Mad Hatter uses a metascore; and our intuition is highly misleading when it comes to metascores.

### What is a metascore

It is important to understand that in general, “metascores” have nothing to do with “metadata”, except for the prefix. This appears to be a common misconception. *Metadata* is data about your experiment: It may be the expected mass accuracy of you measurements, the column type used for chromatography, or the type of sample you are looking at. Such information is already used, to a certain extent, by all existing *in silico* methods: For example, we always have to tell these methods about the expected mass accuracy of our measurements.

In contrast, a “metascore” is simply integrating two or more different scores into one. For example, we may combine the scores of two or more *in silico* methods, such as MetFrag and CFM-ID in [21]. Second, we may combine *in silico* methods and retention time prediction^21–24^. Third, we may combine an *in silico* method with taxonomic information about the analyzed sample^25^. Numerous other types of metascores can be imagined; all of them have in common that they rely on data or metadata of the sample we are analyzing. Such metascores are not what we will talk about in the rest of this papers; they do not have the intrinsic issues discussed in the following.

Yet, in the field of *in silico* methods, the term “metascore” is often used with a different type of method in mind. In many cases, this term describes a method that combines an *in silico* method and “side information” from the structure databases. Mad Hatter is an example of such a metascore, as it uses information from the structure database description (Table 1) to improve its search results. Such metascores do not rely on metadata; instead, they use information that has nothing to do with the sample, the experimental setup or the data.

In the remainder of this manuscript, we will speak about *metascores* when in fact, a more precise denomination would be *“metascore of the* Mad Hatter *type*”. For readability, we will stick to the shorter term. We remind the reader that our critique only applies to those metascores that use “side information” from the searched structure database.

### Blockbuster metabolites

Consider a movie recommendation system that, based on your preferences, suggests new movies to watch. Such recommendation systems widely known these days, in particular from streaming services such as Netflix, Disney+ or Amazon Prime. Yet, a recommendation system may completely ignore the user’s preferences and suggest blockbuster movies all the time: Say, it may choose films according to their financial success. Recommendations will sometimes appear very accurate; after all, we all love blockbuster movies, at least statistically speaking. In evaluations, we will not notice that the recommendation system is basically useless. In practice, we will.

Similar to blockbuster movies, we introduce the term “blockbuster metabolite” for a small molecule that is “better” than others. We stress that this term is meant to include compounds of biological interest, including drugs and toxic compounds. How can a compound be “better” than others? Different features have been suggested to differentiate “interesting” from “uninteresting” compounds: This includes citation counts (the number of papers, patents etc. that contain the compound), production volume, or the number of molecular structure databases a compound is contained in. A metascore will combine the original score that considers the MS/MS data, via some function that also considers, say, the citation count.

A *blockbuster metabolite* is a compound in the structure database we search in, that has a much larger value of the considered feature than all other compounds in the structure database. For example, the compound may have ten thousand citations, whereas most compounds in PubChem have none. In more detail, the compound only needs to have more citations than all compounds it is competing with: *In silico* methods use precursor *m/z* to filter the database for valid candidates, so our compound needs to have a higher citation count than all compounds with a similar mass. The term “blockbuster metabolite” indicates that our metascore may prefer those compounds over all other candidates, independent of the measured MS/MS data.

### Reporting metascore annotation rates is futile

Evaluating a metascore method using a spectral library is meaningless and even misleading. In any such evaluation, we report annotation rates on reference data, with the implicit subtext that somewhat similar annotation rates are reached on biological data. Yet, when evaluating a method that is using a metascore, reported annotation rates have no connection whatsoever with rates that you can reach for biological data. This is a general critique to any such evaluation and does not depend on the utilized metascore.

To understand this phenomenon, consider again our Mad Hatter evaluation above: Mad Hatter correctly annotated about 72% of all compounds when we deliberately removed MS/MS data. The same was possible if we replaced our MS/MS spectra with empty spectra or random spectra, resulting in annotation rates of 71.0% and 71.7%, respectively. It is disturbing that an *in silico* method may basically ignore our data. But the main discrepancy is that we have evaluated a method using a spectral library full of MS/MS spectra, but then ignored all of the data in our evaluation! We could as well have randomly selected a list of compound masses from a structure database; it is very odd to use compounds from a spectral library, which you then plan to ignore. What is the reason to believe that this evaluation will tell us anything about the performance of a method on biological data? We argue that there is no such reason; in fact, there are convincing arguments why we should *not* expect a similar performance for biological data, be it from metabolomics or environmental research.

Now, why is it not a good idea to evaluate a metascore on a spectral library, and what *can* we actually learn from such an evaluation? We can learn that the distribution of compounds in any MS/MS library differs substantially from the distribution of compounds of biological interest, or PubChem compounds. Furthermore, we may learn that it is not very complicated to utilize these differences, to make a metascore shine in evaluation. Yet, the actual number that we record as an “annotation rate”, has no meaning.

For the sake of simplicity, let us focus on citation counts, a feature that has repeatedly been suggested and used for metascores. Similar arguments hold for other features. Fig. 1 shows the distribution of citation counts for different structure databases and libraries. We see that citation counts differ substantially between PubChem, a database containing “molecules of biological interest”, and spectral libraries. We see that most compounds from PubChem have very few citations, whereas biomolecules have larger citation counts. Notably, these are relative abundances: Every compound from the biomolecule structure database is also found in PubChem. Yet, the vast majority of compounds in PubChem are not in the biomolecule structure database, and these compounds usually have very low citation counts. The important point here is that compounds found in a spectral library have many more citations than the average biomolecule, and even more so, than the average PubChem molecule. The reasons for that go in both directions: On the one hand, a compound that has many citations is more likely to be included in a spectral library. On the other hand, a compound that is included in a spectral library will be annotated more often in biological samples and, hence, has a much higher chance to be mentioned in a publication. These are just two reasons, we assume there are many more.

**Fig. 1.**
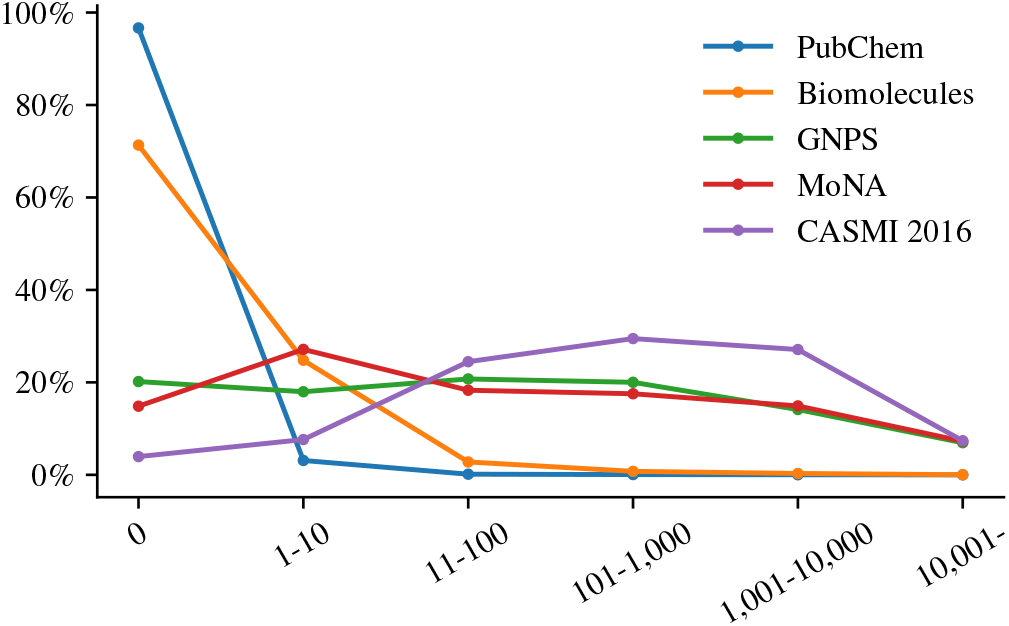
Citation counts. Citations per compound for two structure databases (PubChem, biomolecules) and three spectral libraries (GNPS, MoNA, CASMI 2016 training and testing data). The biomolecule database is a union of several public structure database including HMDB^26^, ChEBI^27^, KEGG^28^ and UNPD^29^. Citation counts are estimated as the number of PubMed IDs associated with the PubChem compound ID, parsed from the file “CID-PMID.gz” provided by PubChem. For the biomolecule database, PubChem, GNPS and MoNA, we uniformly subsampled 10k compounds each. For every compound, we use the maximum citation count of the corresponding molecular structure (first InChIKey block). We excluded rare cases where the molecular structure was not found in PubChem. Dots are connected by lines solely for visualization purpose.

Clearly, there may be compounds in a spectral library that are not blockbuster metabolites: For example, CASMI 2016 contains MS/MS data for 4-[(4-hydroxynaphthalen-1-yl)diazenyl]benzenesulfonic acid, which has only a single citation. These compounds may be wrongly annotated by a metascore as soon as we find a blockbuster metabolite with the same mass. But statistically speaking, such cases are the exception and not the rule: The vast majority of compounds in the CASMI 2016 spectral library are blockbuster metabolites, and using a metascore strongly improves our evaluation results.

Unfortunately, even blind competitions such as CASMI cannot get around this issue, except by explicitly banning the use of metascores. In CASMI 2016, category #3 explicitly allowed metascores such as Mad Hatter. This category was termed “best automatic structural identification — full information”. It explicitly stated that “any form of additional information can be used”, and mentions patents and reference counts as examples. The best submission in this category reached 70% correct annotations. Throughout this paper, we only used information also available to us during the CASMI 2016 contest. Hence, Mad Hatter could have participated in this category, reaching 76% correct annotations and winning the competition. CASMI is a blind competition, and nobody would have even known about our lunatic features.

Notably, not only the evaluation of a metascore method is futile; using annotations from a metascore method to derive statistics about an experimental sample is futile, and even misleading. As discussed, we are mostly annotating blockbuster metabolites and miss out on all other compounds present in the sample. Hence, mapping those annotations to metabolic pathways or compound classes to, say, check for expression change, is not advisable. Doing so will tell us something about blockbuster metabolites and how they map to pathways and compound classes, but very little (if anything) about the data.

### A method shining in evaluations does not have to be reasonable

We have deliberately chosen outrageous features for Mad Hatter. Other metascores often use “reasonable” features such as the number of citations, or the number of biological databases a molecular structure is contained in. The reasoning behind this appears to be as follows: If a metascore sounds reasonable *and* performs well in evaluations, we can safely infer that the approach is justified. Showing that you can build a “good” metascore using outrageous features disproves this implicit reasoning: Mad Hatter works great in evaluations, but it is definitely not a reasonable method. Hence, you must never assume that a method is reasonable or justified simply because it works well in evaluations. A metascore that uses “reasonable” features may or may not work in evaluations; but we cannot infer that the resulting method is reasonable, based on this fact.

Let us take a look behind the curtain: Why is Mad Hatter capable to distinguish between blockbuster and other compounds, despite the fact that the Mad Hatter features are clearly nonsense? For certain features, it is not hard to see why these features favor blockbuster metabolites: For example, blockbusters usually possess a shorter synonym, such as “aspirin” instead of “acetylsalicylic acid” or “2-(Acetyloxy)benzoic acid”. This short synonym is then used as the compound name; consequently, the length of the compound name contains viable information on whether a compound is a blockbuster or not. Another such feature is the number of vowels in the shelf life description: A long description points towards a blockbuster metabolite, and a long description also contains more vowels.

Potentially more dangerous are features for which, even at second thought, we cannot find a reasonable explanation of what is going on. Consider the highly counterintuitive melting point feature: This value is present only for compounds that have been analyzed in some lab, and is left empty for all others. If we had filled in missing values with some reasonable value (say, the average temperature over all compounds where the value is provided), then this feature would carry little useful information for Mad Hatter. Yet, we have taken the liberty to fill in missing values in the worst possible way: We used −273.15°C for those compounds where no melting point was given. This is extremely far off from all compounds that have an entry; hence, this nonchalant little “trick” makes it trivial for machine learning to sort out “uninteresting compounds” where no melting point is provided. Luckily for us, those “uninteresting compounds” are also (almost) never contained in any spectral library. We have hidden the details on missing values somewhere in the methods section, making it hard for a reader to grasp what is going on. We have done this intentionally; yet, we also could have done so out of inexperience with machine learning and how to properly describe it. Recording the temperature in Kelvin plus filling missing values with zeroes is not so far-fetched.

Finally, some features are merely red herrings (say, record number modulo 42, structure complexity); we expect that removing them will not have a large impact on the performance of the metascore, if any. Yet, including them will not impair machine learning, either. Be warned that some of the red herring features may be picked up by machine learning for filtering candidates, in a way unpredictable and possibly incomprehensible for the human brain.

Any type of metascore will, to a varying degree, allow us to annotate blockbuster metabolites and only blockbuster metabolites. It does not really matter if our features are “reasonable” or “unreasonable”; machine learning will find its way to deduce “helpful information” from the data.

### Other compounds become invisible

Blockbuster metabolites are somewhat similar to Hollywood stars: They need all the attention, and they cannot bear rivaling compounds. In detail, blockbuster metabolites will often overshadow other metabolites with the same mass: Irrespective of the actual MS/MS query spectrum that we use for searching, the metascore method will return the blockbuster metabolite. All compounds overshadowed by a blockbuster metabolite become basically invisible. For examples where this goes wrong, see Table 2.

**Table 2.**
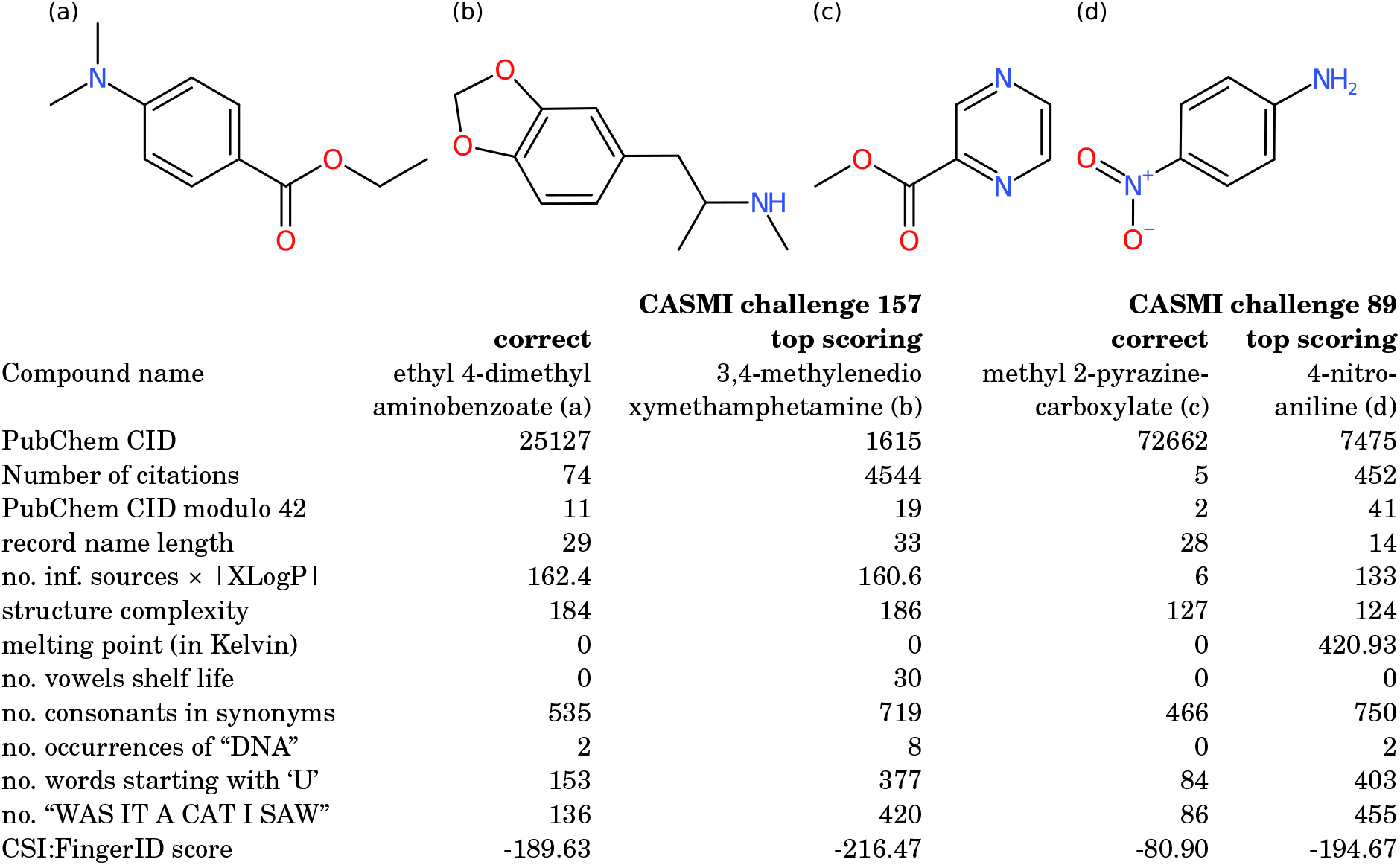
Overshadowing. Left: CASMI 2016 challenge 157. Here, two blockbusters (ethyl 4-dimethylaminobenzoate and 3,4-methylenedioxymethamphetamine) compete with each other, and the bigger blockbuster wins. Right: CASMI 2016 challenge 89. The correct CSI:FingerID answer (methyl 2-pyrazinecarboxylate) is overshadowed by the blockbuster (4-nitroaniline), resulting in an incorrect blockbuster annotation through Mad Hatter.

|  |  |  |  |  |
| --- | --- | --- | --- | --- |
|  | **CASMI challenge 157** |  | **CASMI challenge 89** |  |
|  | **correct** | **top scoring** | **correct** | **top scoring** |
| Compound name | ethyl 4-dimethylaminobenzoate (a) | 3,4-methylenedioxymethamphetamine (b) | methyl 2-pyrazinecarboxylate (c) | 4-nitroaniline (d) |
| PubChem CID | 25127 | 1615 | 72662 | 7475 |
| Number of citations | 74 | 4544 | 5 | 452 |
| PubChem CID modulo 42 | 11 | 19 | 2 | 41 |
| record name length | 29 | 33 | 28 | 14 |
| no. inf. sources × \|XLogP\| | 162.4 | 160.6 | 6 | 133 |
| structure complexity | 184 | 186 | 127 | 124 |
| melting point (in Kelvin) | 0 | 0 | 0 | 420.93 |
| no. vowels shelf life | 0 | 30 | 0 | 0 |
| no. consonants in synonyms | 535 | 719 | 466 | 750 |
| no. occurrences of "DNA" | 2 | 8 | 0 | 2 |
| no. words starting with 'U' | 153 | 377 | 84 | 403 |
| no. "WAS IT A CAT I SAW" | 136 | 420 | 86 | 455 |
| CSI:FingerID score | -189.63 | -216.47 | -80.90 | -194.67 |

Notably, an invisible compound may not appear invisible at first glance: For example, “ethyl 4-dimethylaminobenzoate” (74 citations) was incorrectly annotated as “3,4-methylenedioxymethamphetamine” (4.5k citations, also known as “ecstasy”) in our Mad Hatter evaluation. Clearly, Mad Hatter does not consider citation counts but rather the number of words starting with ‘U’ (153 vs. 377) and the number of “was it a cat I saw?” (136 vs. 420); the result is the same. See Table 2 (left). As noted, blockbuster metabolites are like divas, and the biggest diva takes it all.

Unfortunately, overshadowing is not only an issue when reporting annotation statistics. Assume that you want to annotate methyl 2-pyrazinecarboxylate from Table 2 (right) based on the measured query spectrum. Good news: CSI:FingerID was actually able to annotate the correct compound structure. Bad news: If you are using a metascore, you will never get to know. The metascore replaced the correct answer by an incorrect one (namely, 4-nitroaniline), as the incorrect one was more often cited. Again, it is not citations that Mad Hatter considers, but the result is the same. If we order compound candidates by CSI:FingerID score, we may want to manually correct annotations every now and then, based on citation counts and expert knowledge. Yet, if we only consider the metascore, there is no way for us to spot situations where CSI:FingerID is correct and citation counts are misleading.

The above introduces yet another issue: We might believe that we are searching PubChem when in truth, we are searching in a much smaller database. Using a metascore, we can basically drop all invisible metabolites from the searched structure database without ever changing the search results, regardless of the actual MS/MS query spectrum. Unfortunately, we do not know in which database we are actually searching and, hence, we cannot report this information. This is no good news for reproducibility or good scientific practice. As an example not from CASMI 2016, consider molecular formula C_17_H_21_NO_4_: PubChem contains 8,384 compounds with this molecular formula. Cocaine (PubChem ID 446220) has 35.2k citations, many more than any other compound. It is nevertheless unreasonable to assume that every compound in every sample with this molecular formula is cocaine. Cocaine may even mask other highly cited compounds such as scolopamine (PCID 3000322, 8.9k citations) or fenoterol (PCID 3343, 1.9k citations). But most importantly, more than 8,100 structures in PubChem with this molecular formula do not have a single citation and will be practically invisible for the metascore.

Clearly, there are queries for which overshadowing is not an issue: In particular, there are no blockbusters in the database for certain precursor masses. In these cases, the metascore method will behave like the original method without a metascore. Yet, this does not disprove the above reasoning: For a large fraction of queries, and in particular for biological queries, blockbusters will overshadow other compounds.

### Metabolomics vs. environmental research

So far, we have concentrated on metabolomics and biological data in our argumentation. What about environmental research? There, we have highly reasonable prior information stored in the structure databases: Namely, the production volumes of compounds. The rationale behind this is that something that is not produced by mankind, should also not be annotated in an environmental sample. This appears to be very reasonable, but unfortunately, most of the arguments formulated above remain valid. Clearly, this is true for our argumentation regarding the evaluation setup, discussed in the previous section. But also, with regards to blockbuster compounds, we find that most arguments still hold: An evaluation of a metascore using a spectral library is futile; production rates are a sensible feature to use, but that does not imply that the resulting metascore is sensible; compounds can mask other compounds with lower production volume; excellent hits can be masked by the metascore; we do not really need MS/MS data but instead, could simply use the monoisotopic mass plus the metascore; we cannot report the structure database we are searching in; and so on.

### Targeted vs. untargeted analysis, and the right to refuse to annotate

Studies that employ LC-MS or LC-MS/MS for compound annotation are often referred to as “untargeted” by the authors. This is done to differentiate the instrumental setup from, say, multiple reaction monitoring experiments where we have to choose the compounds to be annotated before we start recording data. Is this nomenclature reasonable? The experimental setup is indeed not targeted to a specific set of compounds.

We argue that doing so is at least misleading, as it mixes up the employed instrumental technology, and the actual experiment we are conducting. Data analysis must be seen as an integral part of any experiment. If we use LC-MS/MS to measure our data, but then limit ourselves to a single compound that we annotate, then this is not an untargeted experiment. Our data would allow us to go untargeted, but if we do not do that, we also should not call it by this name. If we go from one to a thousand compounds, we are still targeting exactly those 1000 compounds. Metaphorically speaking, if you build a propeller engine into a car, you will not fly.

Spectral library search is a “mostly untargeted” data analysis as we restrict ourselves to compounds present in a library, but do not impose further prior knowledge. Any compound in the library has the same chance of getting annotated, as long as it fits the data. A massive advantage in comparison to the “list of masses” is that *we reserve the right not to annotate* certain compounds in the dataset: If the query spectra do not match anything from the spectral library, then we can label those “unknowns”. We are aware that such “unknown” labels are generally considered a pain in metabolomics research^30^. Yet, knowing that we cannot annotate a certain metabolite is definitly better than assigning a bogus annotation!

*In silico* methods are a logical next step for untargeted data analysis, as we extend our search to all compounds in a more comprehensive molecular structure database. Yet, even the structure database is incomplete and we might miss the correct structure. Certain methods try to overcome these restrictions altogether^31–36^, and we expect this to be an active field of research in the years to come.

Using a metascore forces our *in silico* method back into the targeted corset, as invisible compounds can never be found by the metascore. In a way, metascore methods may even be considered “more targeted” than spectral library search, as the later does not give different priors to compounds. Metascores are therefore in fundamental opposition to the central paradigm behind *in silico* methods: These methods were developed to overcome the issue of incomplete spectral libraries and, hence, to make data analysis “less targeted”. It is a puzzling idea to go more and less targeted at the same time. In addition, we mostly lose the possibility to flag certain compounds as “unknowns”, if we search a large database such as PubChem. As we have argued above, the knowledge of what we can and what we cannot confidently annotate, is actually very useful^37^.

## Discussion

This paper is not the first to collect recommendations on how to evaluate machine learning methods in the life sciences^38–40^. It is also not the first to warn about bad evaluations of machine learning models in this area^41^. Notably, Quinn^42^ found that a striking 88% of gut microbiome studies violated established evaluation standards from machine learning. In fact, related warnings were published many decades ago, see for example Ransohoff & Feinstein^43^ and Dreyfus & Dreyfus^44^. This paper is not the first to warn about metascores and blockbuster metabolites^45^, or that more complex machine learning models may perform worse in applications^46^. See also Chapter 12 in [47]. It is not even the first to present its warnings in form of a satire^48^. Yet, by presenting Mad Hatter as a disturbing example of a quirky metascore method reaching excellent evaluation results, we hope to convincingly get the message across that *these points are extremely relevant.*

We believe that editors and reviewers are responsible to end the “reproducibility crisis”^41^ where reported annotation rates of *in silico* methods do not correlate well with annotation rates observed in application. If a computational method is novel, creative and imaginative, there is no need that it outperforms all existing methods to make it publishable. Instead, creating such an artificial barrier forces evaluations into this direction, and holds off authors from evaluating and discussing the limitations of their method. A novel method may well spawn new developments that we cannot even imagine today. Yet, to judge if a computational method is novel, creative and imaginative, you must be an expert in computational methods development. We urge editors to choose qualified reviewers, and we urge reviewers to turn down reviewing invitations if they do not feel fully qualified.

Advocates of metascores have argued that an *in silico* method cannot be used to identify a compound, anyways. If we just want to know which compounds to prioritize for downstream experiments, we may first go after blockbuster metabolites. This is correct, but then, we might as well prioritize compounds by *price.* This would likely result in a highly similar order of candidates, but avoid the false impression that the top-ranked candidate is a better explanation of the recorded data. (In fact, “price” is a confounding variable that may explain many of the observed correlations.) Next, we might want to see what the *in silico* method has to say before going after the blockbuster metabolite annotation: If we are manually analyzing the data anyways, and even consider to order synthetic standards to get it identified, then spending five minutes to look through the result list does not sound excessive. Be warned that using metascore annotations beyond prioritization will inevitably lead to “fake” results: Mapping those annotations to pathways or compound classes will tell us something about blockbuster metabolites and their characteristics, but hardly anything about the data and our sample.

In conclusion, we should accept that an *in silico* method does not have to reach 90% correct annotations to be useful. *In silico* methods cannot do magic; the authors sincerely doubt that MS/MS data will ever allow for such annotation rates, particularly when searching PubChem. In application, we can restrict annotations to, say, a thousand “suspects” and use the precursor *m/z* to annotate them. Doing so, we will wrongly annotate all compounds in our sample that also happen to have one of those masses, to be one of the suspects. Yet, we may decide that this is not a major issue for our analysis. If we decide to measure MS/MS data and want to use an *in silico* method, we should choose a reasonable structure database to search in. PubChem is a wonderful choice for evaluating and comparing methods, but usually a lousy choice in practice. If we notice that certain queries cannot be annotated searching in the smaller structure database with high confidence^37^, we may search those in PubChem. Finally, we may altogether avoid annotating compounds with structures. Structure annotations are not required to, say, compare the distribution of compound classes between samples^32^.

## References

1. Gizzi, A. S., Grove, T. L., Arnold, J. J., et al. A naturally occurring antiviral ribonucleotide encoded by the human genome. Nature 558, 610–614 (2018).

2. Petrick, L. M. & Shomron, N. AI/ML-driven advances in untargeted metabolomics and exposomics for biomedical applications. Cell Rep Phys Sci 3, 100978 (2022).

3. Krettler, C. A. & Thallinger, G. G. A map of mass spectrometry-based in silico fragmentation prediction and compound identification in metabolomics. Brief Bioinform 22, bbab073 (2021).

4. O’Shea, K. & Misra, B. B. Software tools, databases and resources in metabolomics: updates from 2018 to 2019. Metabolomics 16, 36 (2020).

5. Blaženović, I., Kind, T., Ji, J. & Fiehn, O. Software Tools and Approaches for Compound Identification of LC-MS/MS Data in Metabolomics. Metabolites 8, 31 (2018).

6. Hufsky, F. & Böcker, S. Mining molecular structure databases: Identification of small molecules based on fragmentation mass spectrometry data. Mass Spectrom Rev 36, 624–633 (2017).

7. Hufsky, F., Scheubert, K. & Böcker, S. New kids on the block: Novel informatics methods for natural product discovery. Nat Prod Rep 31, 807–817 (2014).

8. Scheubert, K., Hufsky, F. & Böcker, S. Computational Mass Spectrometry for Small Molecules. J Cheminformatics 5, 12 (2013).

9. Schymanski, E. L., Ruttkies, C., Krauss, M., et al. Critical Assessment of Small Molecule Identification 2016: Automated Methods. J Cheminformatics 9, 22 (2017).

10. Nikolić, D., Jones, M., Sumner, L. & Dunn, W. CASMI 2014: Challenges, Solutions and Results. Curr Metabolomics 5 (2017).

11. Nishioka, T., Kasama, T., Kinumi, T., Makabe, H., Matsuda, F., Miura, D., Miyashita, M., Nakamura, T., Tanaka, K. & Yamamoto, A. Winners of CASMI2013: Automated Tools and Challenge Data. Mass Spectrom 3, S0039 (2014).

12. Kim, S., Thiessen, P. A., Bolton, E. E., et al. PubChem Substance and Compound databases. Nucleic Acids Res 44, D1202–D1213 (2016).

13. Srivastava, N., Hinton, G., Krizhevsky, A., Sutskever, I. & Salakhutdinov, R. Dropout: A simple way to prevent neural networks from overfitting. J Mach Learn Res 15, 1929–1958 (2014).

14. Kingma, D. P. & Ba, J. Adam: a method for stochastic optimization 2015. arXiv: 1412.6980.

15. Abadi, M., Barham, P., Chen, J., Chen, Z., Davis, A., Dean, J., Devin, M., Ghemawat, S., Irving, G., Isard, M., et al. TensorFlow: A system for large-scale machine learning in Proc. of USENIX symposium on operating systems design and implementation (OSDI 2016) (2016), 265–283.

16. Dührkop, K., Shen, H., Meusel, M., Rousu, J. & Böcker, S. Searching molecular structure databases with tandem mass spectra using CSI:FingerID. Proc Natl Acad Sci USA 112, 12580–12585 (2015).

17. Dührkop, K., Fleischauer, M., Ludwig, M., Aksenov, A. A., Melnik, A. V., Meusel, M., Dorrestein, P. C., Rousu, J. & Böcker, S. SIRIUS 4: a rapid tool for turning tandem mass spectra into metabolite structure information. Nat Methods 16, 299–302 (2019).

18. Gerlich, M. & Neumann, S. MetFusion: integration of compound identification strategies. J Mass Spectrom 48, 291–298 (2013).

19. Allen, F., Greiner, R. & Wishart, D. Competitive fragmentation modeling of ESI-MS/MS spectra for putative metabolite identification. Metabolomics 11, 98–110 (2015).

20. Ioffe, S. & Szegedy, C. Batch normalization: accelerating deep network training by reducing internal covariate shift in Proc. of International Conference on Machine Learning (ICML 2015) (2015). eprint: 1502.03167.

21. Ruttkies, C., Schymanski, E. L., Wolf, S., Hollender, J. & Neumann, S. MetFrag relaunched: incorporating strategies beyond in silico fragmentation. J Cheminformatics 8, 3 (2016).

22. Menikarachchi, L. C., Cawley, S., Hill, D. W., Hall, L. M., Hall, L., Lai, S., Wilder, J. & Grant, D. F. MolFind: A Software Package Enabling HPLC/MS-Based Identification of Unknown Chemical Structures. Anal Chem 84, 9388–9394 (2012).

23. Bach, E., Szedmak, S., Brouard, C., Böcker, S. & Rousu, J. Liquid-Chromatography Retention Order Prediction for Metabolite Identification. Bioinformatics 34. Proc. of European Conference on Computational Biology (ECCB 2018), i875–i883 (2018).

24. Bach, E., Rogers, S., Williamson, J. & Rousu, J. Probabilistic framework for integration of mass spectrum and retention time information in small molecule identification. Bioinformatics 37, 1724–1731 (2021).

25. Rutz, A., Dounoue-Kubo, M., Ollivier, S., Bisson, J., Bagheri, M., Saesong, T., Ebrahimi, S. N., Ingkaninan, K., Wolfender, J.-L. & Allard, P.-M. Taxonomically Informed Scoring Enhances Confidence in Natural Products Annotation. Front Plant Sci 10, 1329 (2019).

26. Wishart, D. S., Feunang, Y. D., Marcu, A., et al. HMDB 4.0: the human metabolome database for 2018. Nucleic Acids Res 46, D608–D617 (2018).

27. Hastings, J., de Matos, P., Dekker, A., et al. The ChEBI reference database and ontology for biologically relevant chemistry: enhancements for 2013. Nucleic Acids Res 41, D456–D463 (2013).

28. Kanehisa, M., Sato, Y., Kawashima, M., Furumichi, M. & Tanabe, M. KEGG as a reference resource for gene and protein annotation. Nucleic Acids Res 44, D457–D462 (2016).

29. Gu, J., Gui, Y., Chen, L., Yuan, G., Lu, H.-Z. & Xu, X. Use of natural products as chemical library for drug discovery and network pharmacology. PLoS One 8, 1–10 (2013).

30. da Silva, R. R., Dorrestein, P. C. & Quinn, R. A. Illuminating the dark matter in metabolomics. Proc Natl Acad Sci U S A 112, 12549–12550 (2015).

31. Van der Hooft, J. J. J., Wandy, J., Barrett, M. P., Burgess, K. E. V. & Rogers, S. Topic modeling for untargeted substructure exploration in metabolomics. Proc Natl Acad Sci USA 113, 13738–13743 (2016).

32. Dührkop, K., Nothias, L. F., Fleischauer, M., et al. Systematic classification of unknown metabolites using high-resolution fragmentation mass spectra. Nat Biotechnol 39, 462–471 (2021).

33. Litsa, E., Chenthamarakshan, V., Das, P. & Kavraki, L. Spec2Mol: An end-to-end deep learning framework for translating MS/MS Spectra to de-novo molecules. ChemRxiv (2021).

34. Kutuzova, S., Krause, O., McCloskey, D., Nielsen, M. & Igel, C. Multimodal variational autoencoders for semi-supervised learning: In defense of product-of-experts 2021. eprint: 2101.07240.

35. Shrivastava, A. D., Swainston, N., Samanta, S., Roberts, I., Wright Muelas, M. & Kell, D. B. MassGenie: A Transformer-Based Deep Learning Method for Identifying Small Molecules from Their Mass Spectra. Biomolecules 11, 1793 (2021).

36. Stravs, M. A., Dührkop, K., Böcker, S. & Zamboni, N. MSNovelist: de novo structure generation from mass spectra. Nat Methods 19, 865–870 (2022).

37. Hoffmann, M. A., Nothias, L.-F., Ludwig, M., Fleischauer, M., Gentry, E. C., Witting, M., Dorrestein, P. C., Dührkop, K. & Böcker, S. High-confidence structural annotation of metabolites absent from spectral libraries. Nat Biotechnol 40, 411–421 (2022).

38. Chicco, D. Ten quick tips for machine learning in computational biology. BioData mining 10, 35 (2017).

39. Walsh, I., Fishman, D., Garcia-Gasulla, D., Titma, T., Pollastri, G., E.L.I.X.I.R. Machine Learning Focus Group, Harrow, J., Psomopoulos, F. E. & Tosatto, S. C. E. DOME: recommendations for supervised machine learning validation in biology. Nat Methods 18, 1122–1127 (2021).

40. Palmblad, M., Böcker, S., Degroeve, S., Kohlbacher, O., Käll, L., Noble, W. S. & Wilhelm, M. Interpretation of the DOME Recommendations for Machine Learning in Proteomics and Metabolomics. J Proteome Res 21, 1204–1207 (2022).

41. Kapoor, S. & Narayanan, A. Leakage and the Reproducibility Crisis in ML-based Science 2022. arXiv: 2207.07048 [cs. LG].

42. Quinn, T. P. Stool Studies Don’t Pass the Sniff Test: A Systematic Review of Human Gut Microbiome Research Suggests Widespread Misuse of Machine Learning 2021. arXiv: 2107. 03611 [q-bio.GN].

43. Ransohoff, D. F. & Feinstein, A. R. Problems of spectrum and bias in evaluating the efficacy of diagnostic tests. N Engl J Med 299, 926–930 (1978).

44. Dreyfus, H. L. & Dreyfus, S. E. What artificial experts can and cannot do. AI & society 6, 18–26 (1992).

45. Böcker, S. Searching molecular structure databases using tandem MS data: are we there yet? Curr Opin Chem Biol 36, 1–6 (2017).

46. Yaseen, A., Amin, I., Akhter, N., Ben-Hur, A. & Minhas, F. Insights into performance evaluation of compound-protein interaction prediction methods. Bioinformatics 38, ii75–ii81 (2022).

47. Böcker, S. Algorithmic Mass Spectrometry: From molecules to masses and back again. https://bio.informatik.uni-jena.de/textbook-algoms/. Version 0.8.2. Friedrich-Schiller-Universität Jena, Jena, Germany, 2019.

48. Desaire, H. How (Not) to Generate a Highly Predictive Biomarker Panel Using Machine Learning. J Proteome Res 21, 2071–2074 (2022).

